# The EO771 mammary cancer cell line displays a luminal B phenotype and it is sensitive to anti-estrogen treatments

**DOI:** 10.1101/715201

**Authors:** Augustin Le Naour, Yvonne Koffi, Mariane Diab, Delphine Le Guennec, Stéphanie Rougé, Sahar Aldekwer, Nicolas Goncalves-Mendes, Jeremie Talvas, Marie-Chantal Farges, Florence Caldefie-Chezet, Marie-Paule Vasson, Adrien Rossary

**Author notes:** Corresponding author: Augustin Le Naour, *Postal address:* Laboratoire de Biochimie, Biologie moléculaire et Nutrition. Faculté de Pharmacie - 4ème R3. 28 place Henri-Dunant, B.P. 38, 63001 Clermont-Ferrand Cedex 1, *Email address:.

## Abstract

Despite decades of therapeutic trials, detection, many drugs available and numerous studies on breast cancer, it remains the most deadly cancer in women. In order to choose the most appropriate treatment and to know the prognosis of the patients, the breast cancer is divided into different subtypes using a molecular classification. Thus, the need to discover new effective therapy is necessary, requiring to have models to test them. The EO771 (also named E0771 or EO 771) murine mammary cancer cell line is originally isolated from a spontaneous tumor in C57BL/6 mouse. Although frequently used, this cell line remains poorly characterized. Therefore, the EO771 phenotype was investigated. Transcriptomic and protein analysis allowed to classify the EO771 as luminal B subtype and more precisely estrogen receptor α negative, estrogen receptor β positive, progesterone receptor positive and ErbB2 positive. This phenotype was associated to a sensitivity to anti-estrogen treatments such as tamoxifen, 4-hydroxy-tamoxifen, endoxifen and fulvestrant.

## INTRODUCTION

Breast cancer is the most common and deadliest female cancer^1^. The molecular classification of this cancer is now well described. However, preclinical models are needed in order to better understand the development of the cancer and to test new therapies to improve their management. In that way, a mouse syngeneic model of mammary adenocarcinoma can be used by orthotopic injection of EO771 (also named E0771 or EO 771) cells into C57BL/6 mice. This model allows to obtain an immunocompetent model of breast cancer *in vivo*^2,3^. However, this type of mammary tumor remains poorly characterized and the results are divergent concerning its classification. Thus, it is important to better characterize this syngeneic EO771 mammary adenocarcinoma cell line in order to determine the type of tumor that is closest to those found in patients. The expression of hormone receptors such as estrogen receptors alpha (ERα) and beta (ERβ) as well as progesterone receptors (PR) are used routinely by anatomo-pathologists to classify the type of tumor. To complete, the expression of Human Epidermal growth factor receptor 2 (HER2 also named ERBB2). Thus, different tumor subtypes are described: Luminal A characterized by expression of hormone receptors ER+ and/or PR+, absence of overexpression of ERBB2^4^ gene whereas luminal B cancers showed lower expression of ER and PR, but frequently associated to an increased expression of growth factor receptor genes such as HER2^5^. The basal-like or triple negative breast cancer not express any of these markers: ER-, PR-, ERBB2-^4^.

This study first characterized the EO771 cell line concerning its classification and evaluated their sensitivity to anti-estrogen therapy. The identification of signaling pathways, activated after anti-estrogen therapy, was also investigated. Thus, the results shown that EO771 cells displayed a luminal B phenotype, characterized by a phenotype: ERα-, ERβ +, PR+ and ErbB2+. This cell line was sensitive to anti-estrogen drugs such as tamoxifen, which induced an activation of pro-apoptotic signaling pathway as c-Jun NH2-terminal kinase (JNK) and p38 mitogen-activated protein kinase (MAPK) families.

No model of breast cancer derived from C57BL / 6 mice is currently well characterized in terms of their molecular classification. Thus, this work allows to classify this line in the luminal subtype B. These studies are therefore important for transposing the results obtained on this tumor line to patients with luminal B cancer, which corresponds to one of the subtypes most frequently encountered in patients and associated with a poor prognosis.

## MATERIALS AND METHODS

### Mammary adenocarcinoma cell lines

Mouse mammary cancer cell line E0771 (CH3 BioSystems), human triple negative breast cancer cell line MDA-MB-231 (American Type Culture Collection (ATCC), Molsheim, France) and human luminal breast cancer cell line MCF-7 (ATCC) were grown in DMEM supplemented with fetal calf serum (10%), L-Glutamine (1%) and penicillin / streptomycin (1%) at 37°C in 5% CO_2_.

### Cell viability assay

EO771, MCF-7 and MDA-MB-231 cells were plated at a density of 2×10^3^ cells in 96-well plates in a complete medium, and were allowed to adhere in an incubator. Two days later, cells were treated by a range of tamoxifen, 4-OH-tamoxifen, endoxifen or fulvestrant (1 to 25 μM). A supplementation with estradiol (low: 1.5 ng/mL; High: 225 ng/mL), leptin (low: 10 ng/mL; High: 100 ng/mL) or DMSO (0.25%) were also realized. Estradiol concentrations were chosen to be close to the estradiol concentrations measured in mice bearing tumors EO771 observed in our laboratory (data not shown) (low: 1.5 ng/mL; High: 225 ng/mL). Leptin was used at 10 ng/mL (low concentration) and 100 ng/mL (high concentration), close to the physiological conditions^6^ and the final concentration in DMSO was 0.25% in all wells. After 48, 72, and 96 h, cells were washed with PBS and 100 μl of a 25 μg/ml solution of resazurin in DMEM were added to each well. The plates were incubated for 2 h at 37°C in a humidified atmosphere containing 5% CO2. Fluorescence intensity was then measured on an automated 96-well plate reader (Fluoroskan Ascent FL, Thermo Fisher Scientific, Wilmington, DE, USA) using an excitation wavelength of 530 nm and an emission wavelength of 590 nm. Under these conditions, fluorescence was proportional to the number of living cells in the well^7^.

### Treatment of EO771 cells

EO771 cells were plated at a density of 4.5×10^5^ cells in 75cm^2^ flasks in a complete medium, and were allowed to adhere in an incubator. After 48 h, they were treated by the tamoxifen IC50 (14 μM) previously determined. Estradiol (low: 1.5 ng/mL; High: 225 ng/mL), leptin (low: 10 ng/mL; High: 100 ng/mL) and DMSO (final concentration of 0.25% in all flasks) were also added. After 72h of treatments, cells were harvested and protein and RNA were extracted.

### RNA extractions

Total RNA from EO771 cells was extracted by TRIzol^®^ reagent (Invitrogen, Saint Aubin, France) according to the manufacturer’s protocol, and quantified using a NanoDrop spectrophotometer (NanoDrop^®^2000, Thermo Scientific, Waltham, MA, USA). Reverse transcription was performed in a thermocycler (Mastercycler^®^ gradient; Eppendorf, Montesson, France) on 1 μg of total RNA for each condition using a high-capacity cDNA reverse transcription kit (Applied Biosystems, Saint Aubin, France) with random hexamer pdN6 primers.

### Quantitative real-time PCR (q-PCR)

q-PCR was performed using SYBR^®^Green reagents according to the manufacturer’s instructions on a StepOne system (Applied Biosystems). Each condition was assayed in triplicate. Relative quantification was obtained by the comparative Ct method, based on the formula 2^-ΔΔCt^. Expression levels were normalized to the housekeeping gene (GAPDH) for each time point. Sequences and fragment sizes of the human-specific primers used are reported in Table 1.

**Table 1:**
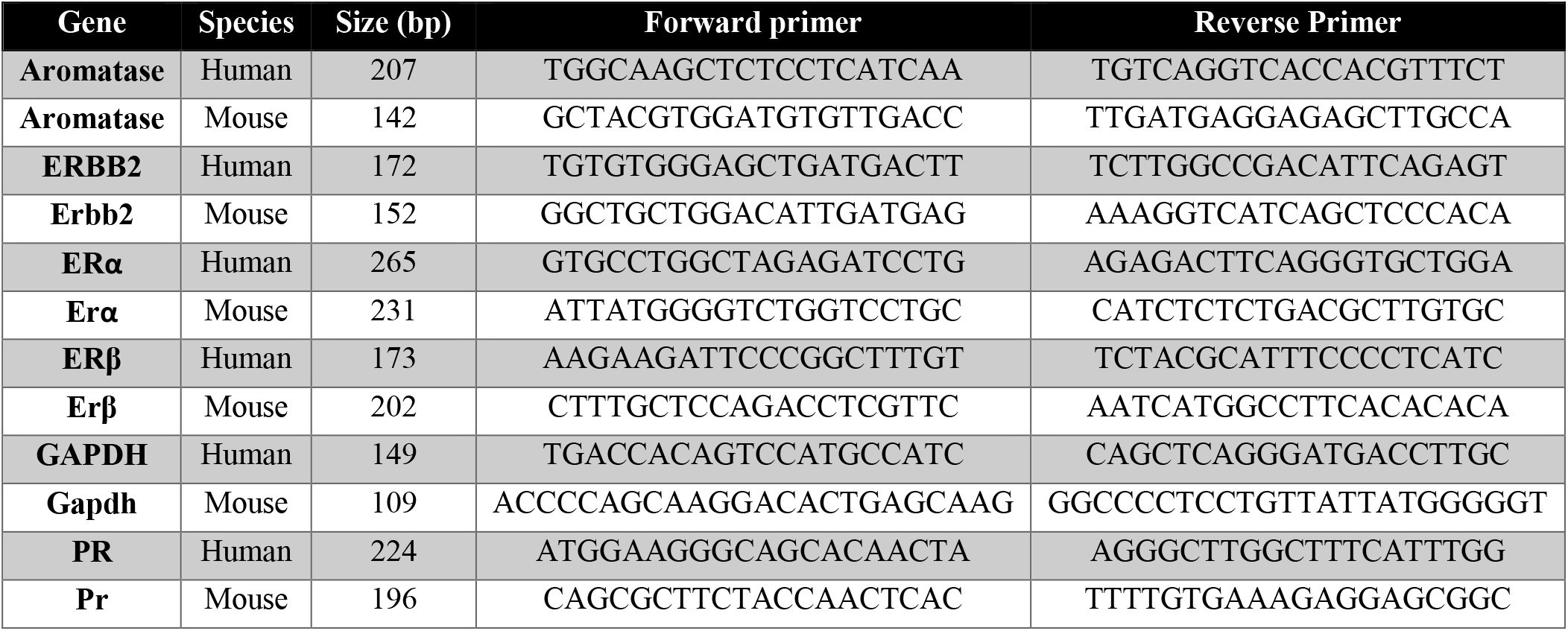
Primers used for qRT-PCR analysis

### Western blot analysis

Protein extractions were performed by RIPA buffer (ThermoFisher Scientific) supplemented by protease (Thermo Scientific Halt Protease Inhibitor Cocktail) and phosphatase (Halt Phosphatase Inhibitor Cocktail) inhibitors. 15 μg of extracted proteins were separated by SDS-PAGE and revealed by antibodies directed against actin (1: 1000, Cell Signaling Technology #8457), GAPDH (1: 1000, Cell Signaling Technology #5174), ERα (1:1000, Abcam #ab32063), ERβ (1:500, Abcam #ab3576), PR (ab133526), ErbB2 (1: 1000, Cell Signaling Technology #4290) and the use of HRP Goat Anti-Rabbit (IgG) secondary antibody (1:5000, Abcam #ab6721).

### Quantification of signaling pathway protein

Using Multiplex Biomarker Immunoassays (cat. kit 48-680MAG and 48-681MA) according to the manufacturer’s instructions, both total and phosphorylated forms of signaling pathways (CREB, JNK, NF-κB, p38, ERK1/2, AKT, p70S6K, STAT3 and STAT5) and MMP3 were determined in EO771 cell line treated or not by the tamoxifen IC50 (14 μM) for 48 hours. The mean fluorescence intensity (MFI) was detected by the Multiplex plate reader for all measurements (Luminex System, Bio-Rad Laboratories, Germany) using a Luminex system, Bio-Rad Laboratories software version 4.2.

### Statistics

For chemoresistance tests, RT-qPCRs and protein analysis the comparison between groups was performed using a Wilcoxon-Mann Whitney test (independent non-parametric data). p-values <0.05 indicate a significant difference. Statistical analyses were performed using GraphPad Prism5 (GraphPad Software, Inc., La Jolla, CA).

## RESULTS

### EO771 cells have a similar phenotype to mammary cancer luminal B subtype

The transcription of genes encoding ERα, ERβ, PR and ERBB2 was evaluated. EO771 cells were compared with human mammary tumor cell line MCF-7 considered to be ER+, PR+, HER2-^8^, ie luminal subtype A, as well as the human mammary tumor cell line MDA-MB-231 admitted as triple negative^9^. Although, the EO771 cells appeared to express ERs (Figure 1A and 1B). They differed to MCF-7 in the transcription of the receptor subtype. Indeed, in the MCF-7 cells, a strong transcription of ERα (Figure 1A) but a small ERβ transcription was observed (Figure 1B). In contrast, a significantly lower transcription of ERα was found in EO771 cells compared to MCF-7 (although its transcription was significantly greater than seen in MDA-MB-231 cells) (Figure 1A). However, the ERβ transcription was significantly greater than observed in MCF-7 and MDA-MB-231 cells (Figure 1B). EO771 cells expressed less PR than MCF-7 cells but this expression was superior compared to MDA-MB-231 triple negative cell line (Figure 1C). The 2 bands observed can be explained by the A and B isoforms of PR, expressed from a single gene^10^. Finally, the EO771 cells did not have an ERBB2 transcription significantly different from the MCF-7 cells, considered not over-expressing ERBB2, but significantly higher than the MDA-MB-231 cells (Figure 1D). In view of these results, the EO771 line could be considered as ERα-, ERβ+, PR+ and ERBB2+/−.

**Figure 1:**
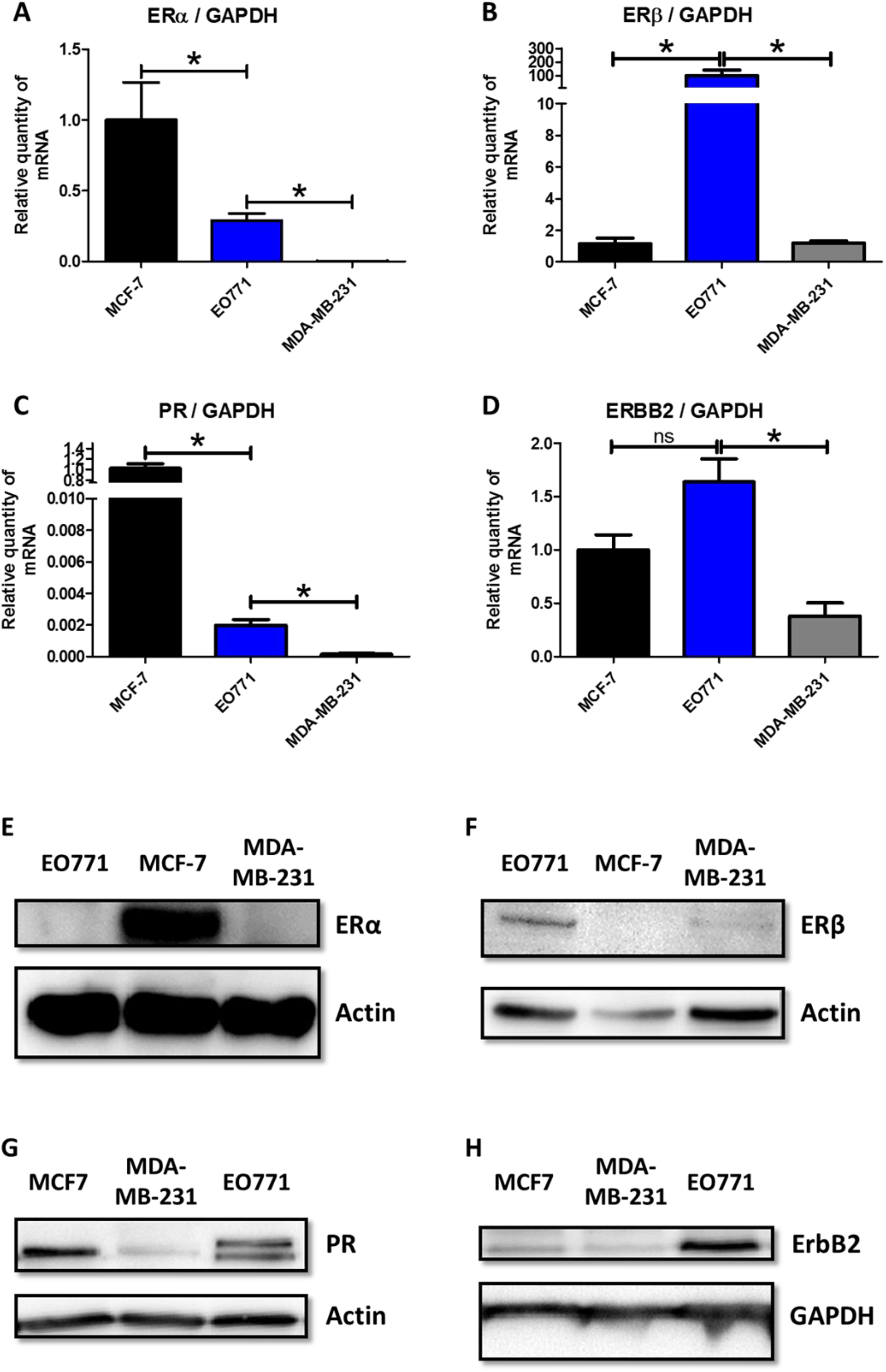
EO771 cells display a luminal B phenotype. A-D: The relative expression of mRNA coding for ERα (A), ERβ (B), PR (C) and ERBB2 (D) was evaluated on MCF-7, MDA-MB-231 and EO771 cells. The values are normalized to the GAPDH gene expression. The data from MCF-7 were set to 1 and the relative quantity of mRNA is shown. P values of <0.05 (*) using a Wilcoxon-Mann Whitney test indicate a significant difference. E-H: ERα (E), ERβ (F), PR (G) and ERBB2 (H) protein levels was assayed by western blot on MCF-7, MDA-MB-231 and EO771 cells (representative of 3 experiments) and normalized to the GAPDH or actin protein levels.

These results were confirmed by evaluating the protein expression of these receptors. Thus, strong ERα expression (Figure 1E) was found for MCF-7 whereas it was undetectable for EO771 and MDA-MB-231 cell line. In contrast, ERβ expression was higher in EO771 cells compared to MCF-7 and MDA-MB-231 cell lines (Figure 1F). The expression of PR in EO771 cells was lower than that observed in MCF-7 (considered as PR+^8^) but superior to MDA-MB-231 (considered as triple negative^9^) (Figure 1G). Finally, concerning the ErbB2 receptor, the expression was greater in the EO771 cells compared with that of the MCF-7 and the MDA-MB-231 (Figure 1H).

Finally, the results of the protein analysis confirm those of the transcriptomic analysis allowing to classify the EO771 as luminal B subtype and more precisely ERα-, ERβ +, PR+ and ErbB2+.

### EO771 cells are sensitive to anti-estrogen treatments

The previous results have shown that EO771 cells expressed few ERα but in return, expressed more ERβ compared to MCF-7 cells. The impact of the expression of these receptors on the sensitivity to anti-estrogenic treatment was evaluated. For that, the sensitivity of EO771 cells was tested to fulvestrant, a competitive estrogen receptor antagonist^11^, and tamoxifen^11,12^ as well as these active metabolites (endoxifen and 4-hydroxy-tamoxifen), all of which are 3 competitive inhibitors of estrogen receptors with partial agonist activity. The MCF-7 cell line, expressing ERα, and the MDA-MB-231, triple negative cell line, were used as controls.

Sensitivity to tamoxifen, 4-hydroxy-tamoxifen (4-OH-tamoxifen), endoxifen and fulvestrant was almost zero in the first 48 hours after treatment in all three cell lines (Figure 2A, 2B, 2C, 2D). After 72 hours of treatment, toxicity of tamoxifen treatments and its active metabolites (endoxifen and 4-OH-tamoxifen) was observed, but only at high dose (15 and 25 μM) on the three mammary cancer cell lines. The sensitivity of EO771 cells was slightly greater with tamoxifen (Figure 2E) and endoxifen (Figure 2G) compared to the two other cell lines. The sensitivity was comparable to that of MCF-7 for 4-OH-tamoxifen (Figure 2F) but these two cell lines were nevertheless more sensitive to these drugs than MDA-MB-231. After 72 hours of treatment with fulvestrant, the viability curves of EO771 and MCF-7 were comparable, showing a greater sensitivity of these two cell lines compared to MDA-MB-231 (Figure 2H). Finally, after 96 hours of treatment, the differences in sensitivity of the three tumor cell lines were more evaluable. As expected, the negative triple line MDA-MB-231 was the least sensitive line of the three, regardless of the hormone therapy used (tamoxifen, 4-OH-tamoxifen, endoxifen and fulvestrant) (Figure 2I, 2J, 2K, 2L). More unexpectedly, a greater sensitivity of the EO771 cells to the four anti-estrogenic molecules was observed, compared to MCF-7 (Figure 2I, 2J, 2K, 2L). Thus, despite a low expression of ERα for EO771 cells compared to MCF-7, it appears that the greater expression of ERβ allows a greater sensitivity to estrogen receptor-targeting treatments.

**Figure 2:**
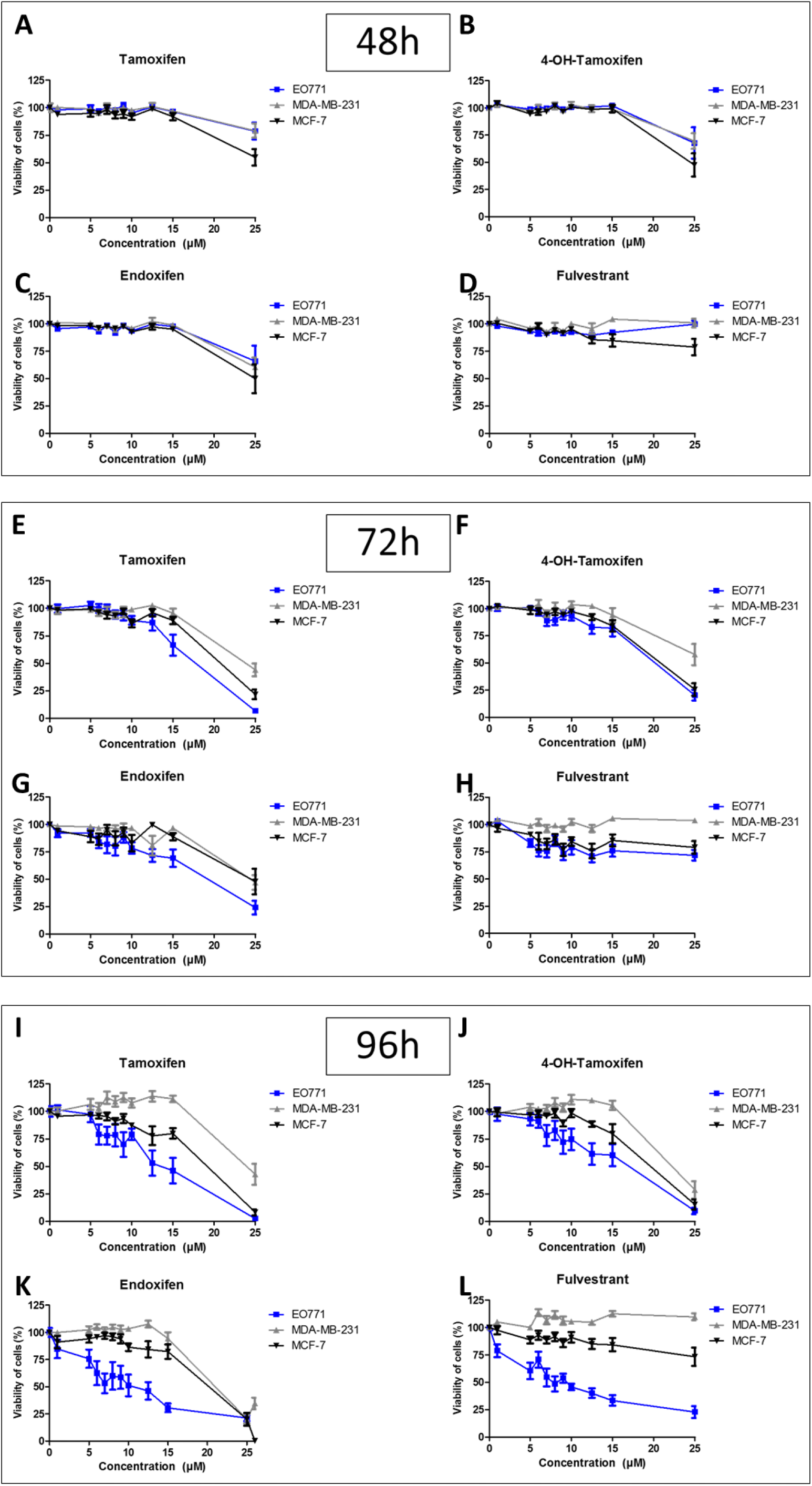
Anti-estrogenic treatments reduce EO771 cells viability. MCF-7, MDA-MB-231 and EO771 cells were treated with increasing tamoxifen (A, E and I), 4-OH-tamoxifen (B, F and J), endoxifen (C, G and K) and fulvestrant (D, H and L) concentrations for 48h (A-D), 72h (E-H) and 96h (I-L). Cell viability was measured by fluorescence using resazurin solution. The untreated condition corresponds to a viability of 100%.

### The treatment with estradiol or leptin does not alter the phenotype neither the sensitivity to tamoxifen of EO771 cells

The presence of estrogens could alter the sensitivity to anti-estrogenic agents in the EO771 cells, inducing competition in their binding to the ERs. Thus, the activity of tamoxifen could be modified in the presence of estradiol (E2). To evaluate this, the expression of hormone receptors in EO771, which would influence their sensitivity to tamoxifen, was investigated in presence of estradiol in the culture medium. The effect of leptin was also evaluated because it decreases tamoxifen activity in MCF-7 cells^13^ by increasing the nuclear expression of ERα^14^ and leptin administration increased plasma estradiol levels^15,16^. The ERα, ERβ, PR expression of the EO771 cells was only slightly modified in the presence of estradiol and leptin whether in the absence (Figure 3A) or in the presence of tamoxifen (Figure 3B).

**Figure 3:**
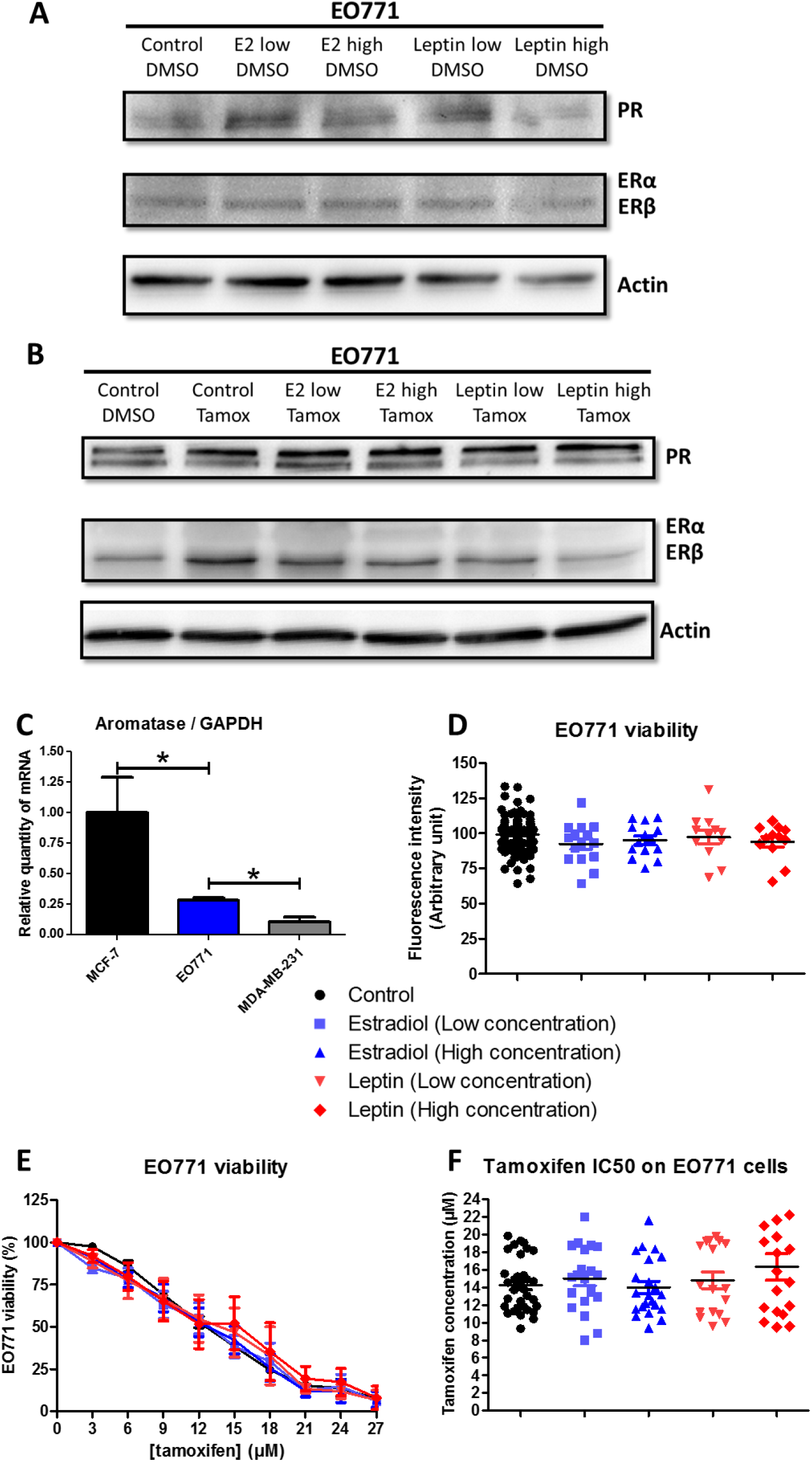
Estradiol and leptin do not influence EO771 cells phenotype and viability. A, B: ERα, ERβ and PR protein levels was assayed by western blot on EO771 cells (representative of 3 experiments) and normalized to the actin protein levels in presence of estradiol (Low: 1.5 ng/mL; High: 225 ng/mL) or leptin (Low: 10 ng/mL; High: 100 ng/mL) in untreated condition (DMSO) (A) or with tamoxifen (B). C: The relative expression of mRNA coding for aromatase was evaluated on MCF-7, MDA-MB-231 and EO771 cells. The values are normalized to the GAPDH gene expression. The data from MCF-7 were set to 1 and the relative quantity of mRNA is shown. P values of <0.05 (*) using a Wilcoxon-Mann Whitney test indicate a significant difference. D-F: Cell viability was measured by fluorescence using resazurin solution. EO771 cells were cultured in the presence of estradiol, leptin or vehicle (D) and EO771 cells were treated with increasing tamoxifen concentration (E). The untreated condition corresponded to a viability of 100%. The tamoxifen IC50 corresponded to the tamoxifen concentration inducing 50% of EO771 viability (F).

Before, to evaluate the sensitivity of EO771 cells in the presence of estradiol or leptin, the measure of the expression of the gene coding for aromatase (enzyme allowing the production of estrogens by androgen transformation) was studied in order to evaluate whether these cells were significantly able to produce estradiol. The gene coding for aromatase was very poorly transcribed in EO771 cells (Figure 3C). The study of protein expression by western blot did not detect aromatase in EO771 cells (data not shown). Thus, these cells did not seem able of producing large amounts of estradiol.

In our experimental conditions, the addition of estradiol or leptin did not modify the tumor growth of EO771 cells (Figure 3D). Likewise, the sensitivity of EO771 cells to tamoxifen treatment was not modified with respect to the control condition (Figure 3E). That was supported by the calculation of the concentrations of tamoxifen inhibiting 50% of cell viability (IC50). In fact, the IC50 values of tamoxifen for EO771 cells in the presence of estradiol or leptin were not significantly different compared to the control condition (Figure 3F).

Thus, the presence of estradiol and leptin did not alter the luminal B phenotype nor the sensitivity to tamoxifen of EO771 cells.

### Tamoxifen activates signaling pathways in EO771 cells

Anti-estrogenic treatment leads to a cytotoxic effect on EO771 cells. However, the presence of estrogen has no effect on either proliferation, sensitivity to tamoxifen, or expression of hormone receptors. Thus, the response to tamoxifen observed in EO771 cells may be independent of estrogen receptors. Then, the signaling pathways, which could be affected by tamoxifen treatment, were studied.

The mitogen-activated protein kinase (MAPKs) family pathway, which is a family of kinases that transduce signals from the cell membrane to the nucleus in response to numerous stimuli, was investigated. Classically, MAPKs are divided into three extracellular signal-regulated kinase (ERK) families that have an anti-apoptotic role, and the c-Jun NH2-terminal kinase (JNK) and p38-MAPK families which are both associated to stress pathways leading to a pro-apoptotic effect^17^. The presence of tamoxifen led to a pro-apoptotic signal in EO771 cells by activating the JNK and p38-MAPK pathways (Figure 4A and 4B) without activating the ERK anti-apoptotic pathway (Figure 4C).

The nuclear factor-κB (NF-κB) pathway has been shown to be inhibited by tamoxifen^18^. Thus, in our experiment, the presence of tamoxifen did not modify the NF-κB pathway activation in EO771 cells (Figure 4D).

**Figure 4:**
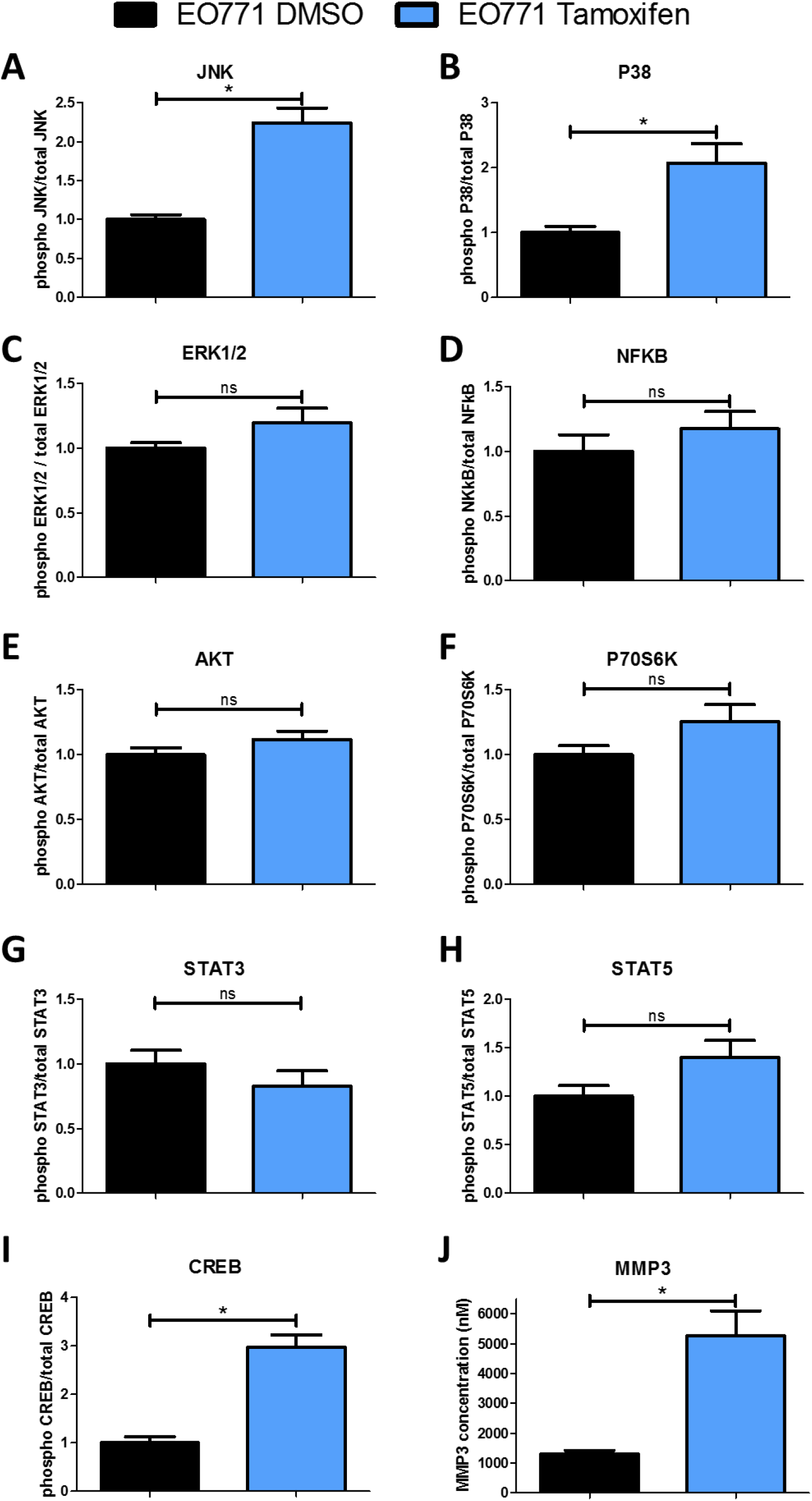
Tamoxifen modify intracellular signaling pathways in EO771 cells. EO771 cells were treated by the tamoxifen IC50 (14μM) or vehicle (DMSO) for 48 hours. The expression of both total and phosphorylated forms of signaling pathway proteins (CREB, JNK, NFκB, p38, ERK1/2, AKT, p70S6K, STAT3 and STAT5) and MMP3 was analyzed by measure of fluorescence intensity (MFI) using a Luminex system. P values of <0.05 (*) using a Wilcoxon-Mann Whitney test indicate a significant difference.

The phosphoinositide 3 kinase (PI3K)/AKT pathway is a survival pathway leading to enhanced cell survival and cell cycle progression. In EO771 cells, the addition of tamoxifen did not cause any change in the activation of this pathway (Figure 4E). Similarly, the p70S6K kinase, which is phosphorylated and activated by mTOR in mitogenic pathways downstream of PI3K/AKT, was not differently phosphorylated in tamoxifen-treated EO771 cells compared to the control condition (Figure 4F).

Signaling pathways involving the Signal Transducer and Activator of Transcription (STAT) 3 and STAT5 were also studied. Indeed, STAT5 assumes essential roles in proliferation, differentiation and survival of multipotent mammary stem cells^19^. STAT3 and STAT5 expression can be found in all breast cancer subtypes and a down-regulation of both by different drugs was associated with reduced growth in breast cancer subtype^20^. In our model, tamoxifen did not significantly affect the activation of STAT3 and STAT5 pathways compared to the control condition (Figure 4G and 4H). Thus, these pathways did not appear to be involved in the cytotoxic activity of tamoxifen on EO771 cells.

Chen *et al.* have shown that downregulation of cAMP-response element binding protein (CREB) was associated with inhibition of mammary tumor cell growth by a mechanism that appears to be independent of ER as acting on both triple negative cells (MDA-MB-231) and ER+ cells (MCF-7)^21^. Knowing that this pathway is found activated during resistance to tamoxifen^22^, the effects of this drug on CREB pathway were investigated in EO771 cells. Interestingly, this pathway was found significantly increased during treatment with tamoxifen (Figure 4I).

Finally, the potential effect of tamoxifen on metastic dissemination of EO771 cell was investigated. For this, the production of matrix metalloproteinase-3 (MMP-3), which might be involved in metastatic dissemination of breast cancer^23^, was evaluated. An increase in MMP3 production in tamoxifen-treated EO771 cells was observed, compared to untreated cells (Figure 4J). Thus, despite the cytotoxic activity of tamoxifen on EO771 cells, this treatment could promote metastatic spread due to increased production of MMP3.

In summary, EO771 presented a luminal B phenotype with the expression of ERβ, PR and ErbB2. They were sensitive to tamoxifen, which induced activation of pro-apoptosis pathways, such as p38 and JNK but also an increased the MMP3 level (Figure 5).

**Figure 5:**
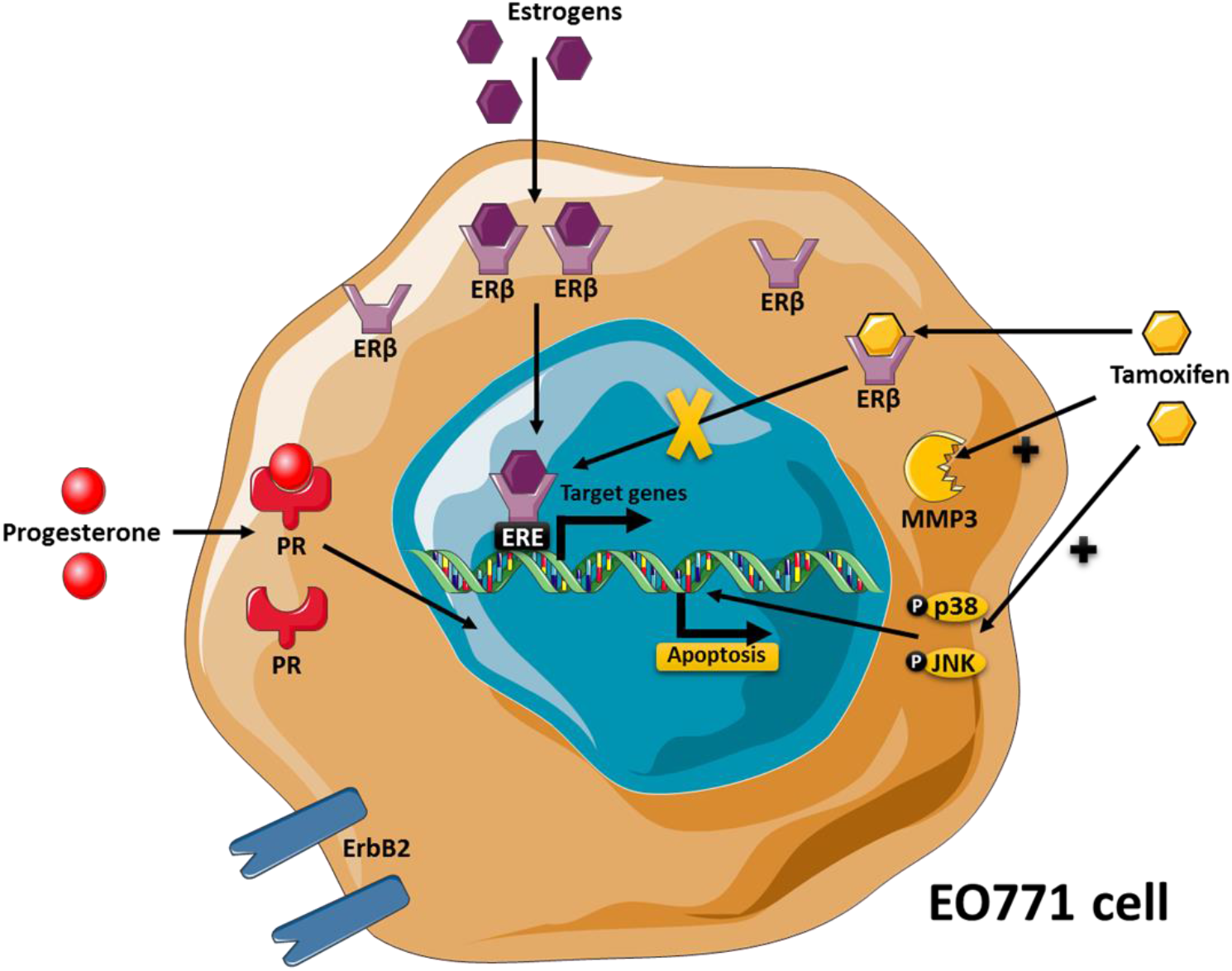
The EO771 phenotype and their sensitivity to tamoxifen. EO771 cells express ERβ, PR and HER2. In presence of estrogens, ERβ are translocated to the nucleus where they bind to the estrogen response elements (ERE) inducing transduction of target genes. Anti-estrogen treatments, such as tamoxifen, block this effect. On the contrary, tamoxifen induce the activation of p38 and JNK pathway, leading to apoptosis, but also increase the MMP3 level.

## DISCUSSION

The need to find mouse models to mimic breast cancer is essential to improve the management of this cancer, which remains the deadliest in women. EO771 cells, derived from a spontaneous tumor of C57BL/6 mice^24^, are used in syngeneic models^3,14^. This model, although frequently used, remained poorly characterized. This work shows that EO771 cells expressed hormonal receptors, and more particularly ERβ. The phenotype study of these cells also showed that they expressed HER2 more than MCF-7 cells. Thus, these cells exhibited the characteristics of luminal B subtype^5^, ER+, PR+ and HER2+. According to the literature^25^, a lower expression for ERα and PR was observed compared to luminal A cells (MCF-7). These frequent tumors are found in about 30 to 40% of breast cancers^25,26^ and are generally more aggressive, with high grade, and have a worse prognosis than luminal A breast cancers^5^. As in patients with luminal B tumor, EO771 cells were sensitive to tamoxifen therapy suggesting that ERβ would mediate its anti-tumor activity, whereas it is generally associated with ERα. This line could also be derived from breast cancer initiating cells because Ma *et al*. have reported an absence of ERα but an upregulation of ERβ in breast cancer cells with tumor-initiating capabilities with phenotypic stem cell markers^27^. Thus, knowing now the phenotype of this line EO771, the latter can be used to test many anti-tumor molecules^5^ such as selective inhibitors of ERβ^28^.

Among the molecules conventionally used to treat luminal cancers, tamoxifen induced in these cells the activation of pro-apoptotic pathways involving JNK and p38-MAPK. These latter exert an anti-tumoral effect interesting to treat for treating this type of tumor. However, the use of this drug was also associated with an increase in MMP3, known to possess pro-metastatic activity. Indeed, this MMP3 was found to be increased in EO771.LMB cells, isolated from a spontaneous lung metastasis from an EO771 tumor-bearing compared with parental EO771^23^. It would be interesting to see if treatment with tamoxifen would not induce an increase in metastatic spread in a mouse model with EO771 mammary tumors.

Thus, the phenotyping of the EO771 line classified this line in the luminal subtype B allowing a parallel between the results of the *in vitro* and *in vivo* studies, obtained with this murine model, and luminal B breast cancers encountered in patients. This EO771 cell line corresponds to one of the subtypes most frequently encountered in patients and associated with a poor prognosis.

## AKNOWLEDGEMENTS

This work was supported by funding from l’Institut National du Cancer (INCA: MammAdipo project; PLBIO 13-106) and Le comité de l’Allier de la Ligue contre le cancer.

